# The spatiotemporal link of temporal expectations: contextual temporal expectation is independent of spatial attention

**DOI:** 10.1101/2021.07.30.454407

**Authors:** Noam Tal-Perry, Shlomit Yuval-Greenberg

## Abstract

Temporal expectation is the ability to construct predictions regarding the timing of events, based on previously-experienced temporal regularities of different types. For example, cue-based expectations are constructed when a cue validly indicates when a target is expected to occur. However, in the absence of such cues, expectations can be constructed based on contextual temporal information, including the event’s hazard-rate function – its moment-by-moment conditional probability that changes over time; and prior experiences, which provide probabilistic information regarding the event’s predicted timing (sequential effects).

It was previously suggested that cue-based temporal expectation is exerted via synchronization of spatially-specific neural activity at a target's predictable time, within receptive fields corresponding to the target’s expected location. Here, we tested if the same theoretical model holds for contextual temporal effects. Participants (n = 40) performed a speeded spatial-cueing detection task, with two-thirds valid spatial cues. The target’s hazard-rate function was modulated by varying the foreperiod – the interval between the spatial cue and the target - among trials, and was manipulated between groups by changing the interval distribution. Reaction times were analyzed using both frequentist and Bayesian generalized linear mixed models, accounting for hazard and sequential effects. Results showed that the effects of contextual temporal structures on reaction times were independent of spatial attention. This suggests that the spatiotemporal mechanisms, thought to account for cue-based expectation, cannot explain other sources of temporal expectations. We conclude that expectations based on contextual structures have different characteristics than cue-based temporal expectation, suggesting reliance on distinct neural mechanisms.

**Significance statement:** Temporal expectation is the ability to predict an event onset based on temporal regularities. A neurophysiological model suggested that temporal expectation relies on the synchronization of spatially-specific neurons whose receptive fields represent the attended location. This model predicts that temporal expectation would be evident solely within the locus of spatial attention. Existing evidence supported this model for expectation based on associations between a temporal cue and a target, but here we show that it cannot account for another source of temporal expectation – expectation that is based on contextual information, i.e. hazard-rate and recent priors. These findings reveal the existence of different predictive mechanisms for cued and contextual temporal predictions, with the former depending on spatial attention and the latter non-spatially-specific.

## Introduction

Temporal expectation is the ability to construct predictions regarding the timing of events, based on temporal regularities. Multiple forms of such regularities can drive temporal expectation, including contextual information, when information regarding distributions of events and statistical inferences from recent experiences are used to predict the timings of future events; rhythms and other repetitive sequences, when events occur in predictable streams (e.g., Heideman et al., 2016; Breska and Deouell, 2017; Dankner et al., 2017; Breska and Ivry, 2018); and cued-associations, when events are preceded by informative temporal cues (e.g., Coull and Nobre, 1998; Miniussi et al., 1999). Studies show that expectations of all these sources are associated with enhanced perceptual performances (e.g., Niemi and Näätänen, 1981; Nobre et al., 2007; Nobre and van Ede, 2018).

Despite abundant evidence on behavioral effects of temporal expectation, relatively little is known regarding their neurophysiological correlates. One theoretical framework suggested that temporal expectation is the result of synchronization within neural populations at the time of the expected target. It was suggested that these neuronal populations are spatially specific – their receptive fields correspond to the expected target location (Rohenkohl et al., 2014; Nobre and van Ede, 2018). According to this view, temporal and spatial expectations are tightly linked, as temporal expectation is bound to be evident only within the locus of spatial attention: in order to gain from knowing *when* a target will occur, one has to know *where* it would occur. However, evidence for this spatiotemporal framework is limited to studies that manipulated cue-based temporal expectation (Doherty et al., 2005; Rohenkohl et al., 2014; Seibold et al., 2020). It remains unknown whether the same spatiotemporal mechanism accounts for temporal expectation based on other sources of regularities. Here, we examine whether this spatiotemporal framework could also explain expectations based on contextual information, i.e., induced by conditional probabilities or sequential effects.

C*onditional probability* is the likelihood of an event to occur, given that it has yet to occur. This probability changes continuously as time progresses and can be described as a function of time, termed the *hazard-rate* function. When the timings of events are uniformly distributed, the hazard-rate function is monotonically increasing, but other distributions would lead to different hazard-rate functions (Luce, 1986). The effect of the hazard-rate function was demonstrated by showing that higher conditional probability for target occurrence is associated with enhanced performance. In a common design, a warning signal (WS) alerts participants to an upcoming target, which follows after a varying time-interval (*foreperiod*). It is consistently found that performance for targets appearing following long foreperiods is enhanced relative to targets appearing following shorter ones (Näätänen, 1970; Niemi and Näätänen, 1981).

Another source of information used to alleviate temporal uncertainty are prior experiences. The perceptual system constantly makes predictions and utilizes priors to make these predictions (Clark, 2013). These temporal predictions about the event's most probable onset time are reflected in the *sequential effect* – the cost and benefit in performance stemming from the relation between the foreperiods of sequential trials (Bertelson, 1961; Niemi and Näätänen, 1981). When a target appears following a foreperiod that is shorter than that of the previous trial, performance is reduced, relative to trials that were preceded by an identical foreperiod. This pattern is asymmetrical, as performance remains unchanged when a target appears following a foreperiod that is longer than the previous trial (Bertelson, 1961; Possamai et al., 1973).

Here, we manipulated spatial attention and temporal expectation simultaneously. In each trial, participants were presented with a spatial cue that was either congruent, incongruent, or neutral in respect to the location of the target that appeared after a varying interval (foreperiod). The distribution of the foreperiod intervals was varied between participants to create two different hazard-rate functions. We hypothesized that, unlike cue-based expectation, both hazard-rate and sequential effects are independent of spatial attention, indicating that the spatiotemporal framework suggested to account for temporal expectation does not account for these processes.

## Materials and Methods

### Participants

A total of 40 participants were included in this study, 20 in the ‘Uniform distribution’ group (12 females, 2 left-handed, Mean age 25.35±3.5 standard deviations [SD]) and 20 in the ‘Inverse U-shape distribution’ group (13 females, one left-handed, Mean age 24.55±4.0 SD). Participants received payment or course credit for their participation. All participants were healthy, reported normal or corrected-to-normal vision, and no history of neurological disorders. The experimental protocols were approved by the ethical committees of Tel-Aviv University and the School of Psychological Sciences. Prior to participation, participants signed informed consent forms.

### Stimuli

The fixation object consisted of a dot (0.075° radius) within a ring (0.15° radius), embedded within a diamond shape (0.4×0.4°). The edges of the diamond changed color from black to white, cueing attention to the left (two left edges became white) or right (two right edges became white) side of fixation object, or remaining neutral in respect to target location (all four edges became white) (**see Fig. 1**). The target was a black asterisk (0.4×0.4°) presented at 4° eccentricity to the right or left of fixation object. A 1000 Hz pure tone was sounded for 60 ms as negative feedback following errors. Fixation object and target were presented on a mid-gray background.

**Figure 1.**
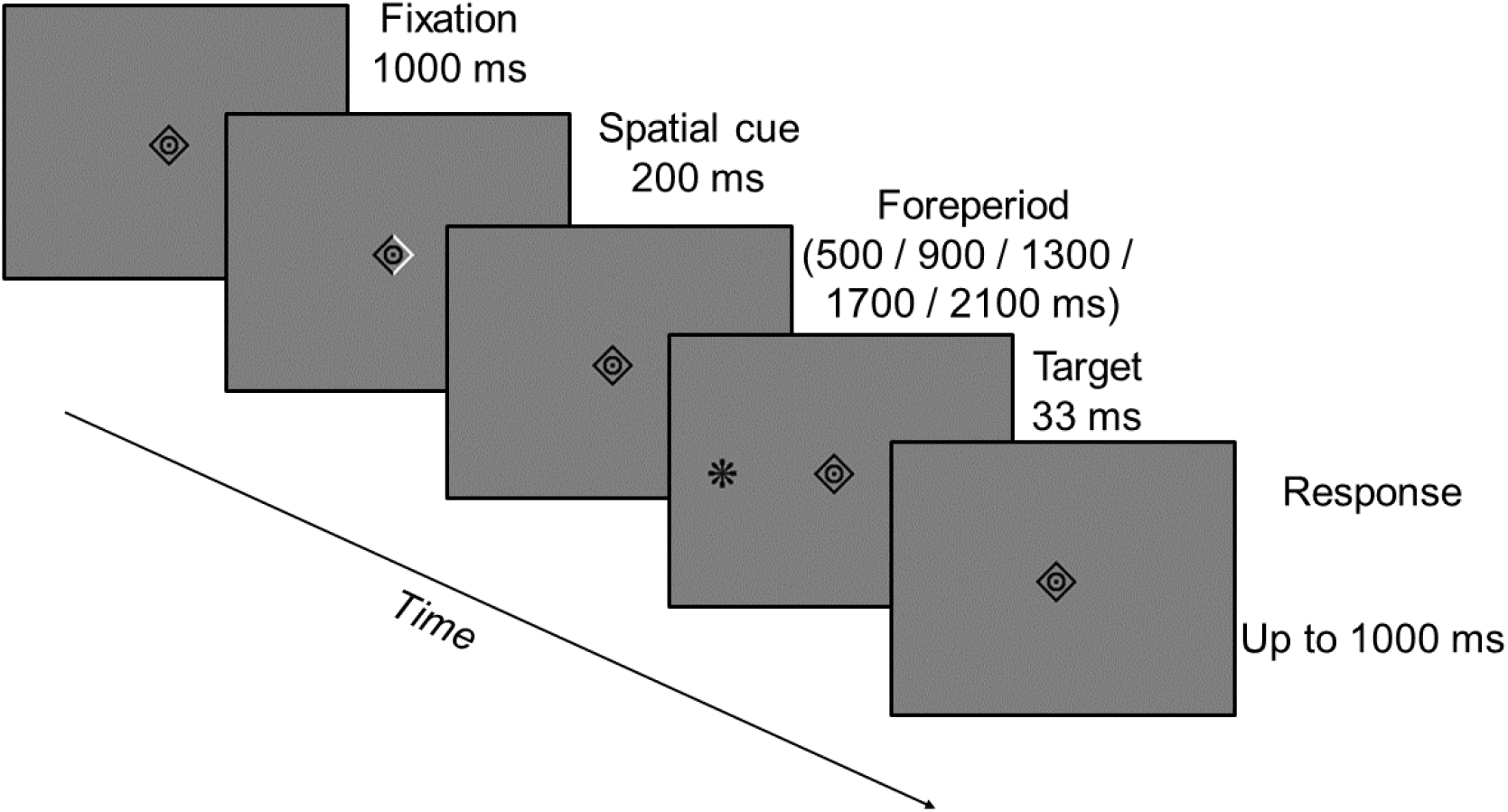
*Trial progression*. Fixation period lasted until stable fixation was confirmed with online eye tracking procedure. Spatial cue was invalid in respect to target location in 25% of trials (as depicted), valid in 50% of trials and uninformative trials in 25% of trials. In two groups, foreperiods were sampled from either a uniform or an inverse-U distribution. Stable fixation was enforced during the foreperiod using online gaze-contingency. Participants were asked to make a single-button speeded response within 1000 ms of target onset. An error tone was played when participants responded before target onset, or failed to respond within the time limit. Stimuli size and eccentricity increased for display purposes and are not to scale

### Experimental design

Participants were seated in a dimly lit room, with a computer monitor placed 100 cm in front of them (24" LCD ASUS VG248QE, 1,920 × 1,080 pixels resolution, 120 Hz refresh rate, mid-gray luminance was measured to be 110 cd/m^2^). During the experiment, participants rested their heads on a chinrest. MATLAB R2015a (Mathworks, USA) was used to code and control the experiment, with stimuli displayed using Psychophysics Toolbox v3 (Brainard, 1997). Gaze position was monitored binocularly using EyeLink 1000 Plus infrared video-oculographic desktop mounted system (SR Research Ltd., Oakville, ON, Canada) throughout the experiment, at a sampling rate of 1000 Hz. This system has <0.01° spatial resolution and an average accuracy of 0.25–0.5° when a chinrest is used, according to the manufacturer. A nine-point calibration of the eye-tracker was performed prior to each block and whenever necessary.

Each trial started with a central black fixation object, presented until an online gaze-contingent procedure verified 1000 ms of stable fixation (gaze was placed within a radius of 1.5° of screen center). Following this, the edges of the fixation object changed color for 200 ms to represent a spatial informative or uninformative cue. After a varying foreperiod (500 / 900 / 1300 / 1700 / 2100 ms) the target was briefly (33 ms) presented at 4° to the left or right of center, with target being congruent to a spatially-informative cue direction in 50% of trials (valid condition), incongruent in 25% of trials (invalid condition), or neutral with respect to a spatially-uninformative cue in the remaining 25% of trials (uninformative condition). Participants were requested to press a key with their dominant hand, as quickly as possible and after no longer than 1000 ms, upon target detection. Between groups, participants were presented with the five foreperiods in either a uniform distribution (20% probability for each foreperiod) or an inverse-U-shaped distribution (a ratio of 1:2:3:2:1 between the five foreperiods, leading to trial percentages of approximately 11%, 22%, 33%, 22%, and 11%, respectively). These prior distributions resulted in different time-dependent conditional probabilities, i.e. different hazard-rate functions, as depicted in **Fig. 2**. The manipulation of hazard-rate was required to differentiate its effect from other foreperiod effects related to the WS, such as arousal (Steinborn and Langner, 2012; Weinbach and Henik, 2012). The different distributions were examined in separate participant groups, in order to avoid carry-over effects of distribution learning (Mattiesing et al., 2017). Fixation was monitored throughout the foreperiod, using an online gaze-contingent procedure, and trials that included ≥ 1.5° gaze-shift for more than 10 ms during this period were aborted and repeated at a later stage of the session. An error feedback tone was sounded when participants responded before target onset or did not respond within 1000 ms following target onset. These trials were not included in the analysis. The trial procedure is depicted in **Fig. 1**.

**Figure 2.**
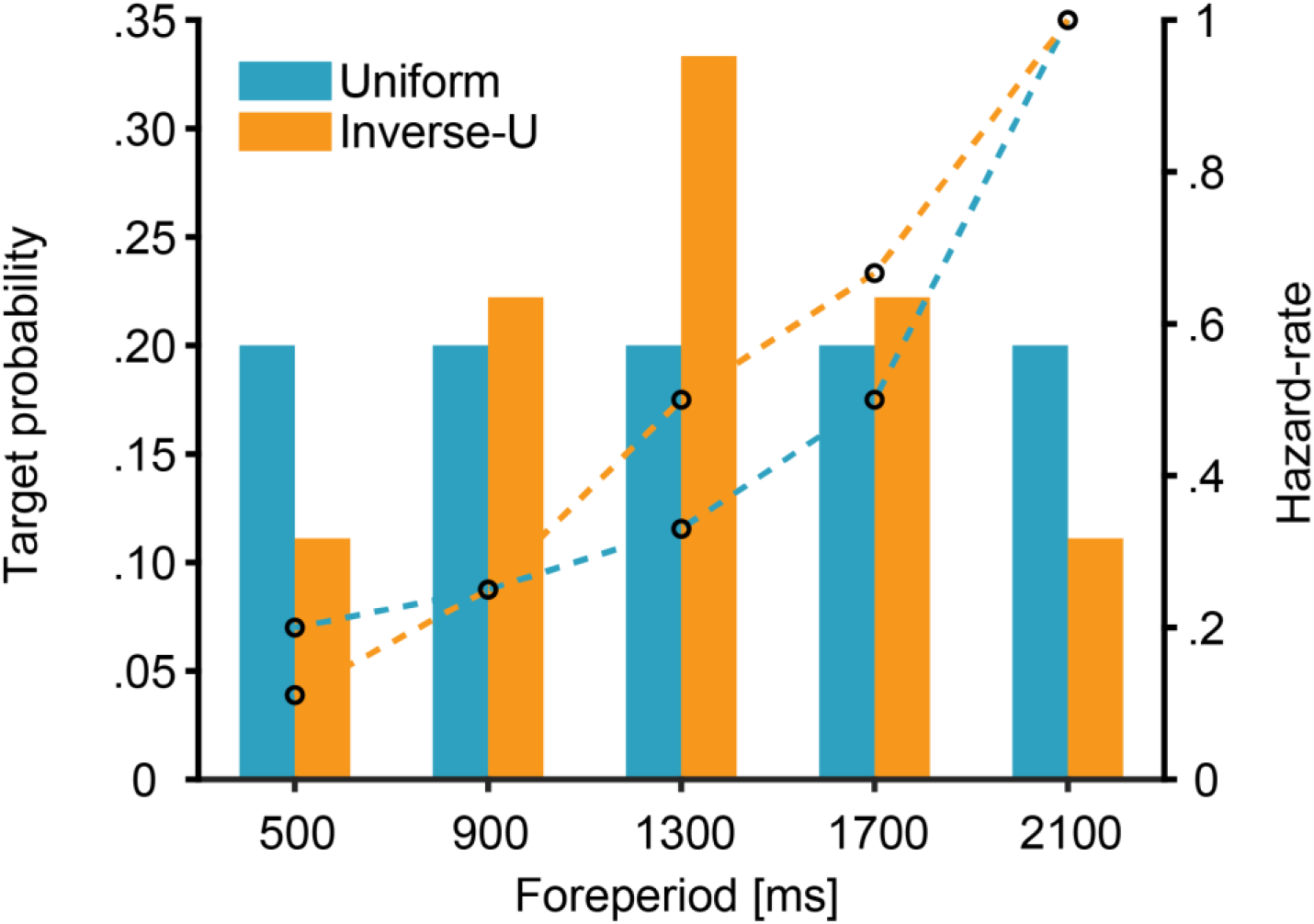
Target probability (bars) and hazard-rate (conditional probability, dashed line) for the uniform and inverse-U foreperiod distributions

Participants of the uniform distribution group performed 10 blocks of 160 trials each, divided into two sessions of approximately 1.25 hours each. Participants of the inverse-U-shaped distribution group performed 18 blocks of 144 trials each, divided into three sessions of approximately 1.25 hours each. This number of repetitions guaranteed that we have a minimum of 50 trials in all conditions and for all foreperiods in each of the two distributions, and a large enough number of trials conduct a sequential analysis on pairs of consecutive trials. A short break was given after each block. Feedback on performance in each block was provided at the end of each experimental block and included: mean RT and number of error trials (including both missed trials or premature responses). Starting from the 2^nd^ experimental block, participants were also presented with a message that encouraged them to perform faster if the current block's mean RT fell below their global mean RT of the entire session. A practice block of 10 trials with random conditions was administered at the beginning of each session.

### Statistical analysis

A negligible amount of trials with no response (< 1% of all trials; mean 0.7% of trials per participant, range 0-2.16% of trials) were discarded from analysis. Additionally, trials with response time below 150 ms were considered unlikely to represent genuine target-related responses (Keele and Posner, 1968; McLeod, 1987) and were likewise discarded from analysis (< 1% of all remaining trials; mean 0.3% of trials per participant, range 0-2.2% of trials).

The reaction times (RTs) of the remaining trials were modeled using a generalized linear mixed model (GLMM), assuming a gamma family of responses with an identity link (see explanation below) (Baayen and Milin, 2010; Lo and Andrews, 2015). Unlike analysis of variance (ANOVA), GLMM is suited for non-normally distributed variables, like the positively skewed RT distribution, while also allowing to model trial-level covariates, thus increasing the analysis’ power (Baayen and Milin, 2010). Hierarchical models are also well suited for unevenly distributed trial numbers among conditions, as is the case with the Inverse-U shaped distribution and the sequential effect in the current study, by weighting the population-level mean according to the number of samples included in the subject-level means for each condition. An assumption of this analysis is that the RTs follow Gamma distribution. Gamma distributions are suited to describe continuous responses that are zero-bounded and have a unimodal and rightward-skewed distribution (e.g., RTs). We further assumed that the predictors are linearly related to the predicted RT, thus an identity link was used (i.e., no transformation was made on the value produced by the predictors) (Lo and Andrews, 2015).

The following fixed effects were modeled: (1) linear and quadratic terms for Foreperiod duration, to model the slope of the foreperiod effect; (2) Cue (valid / invalid / uninformative), to model the effect of spatial attention; (3) the Foreperiod (FP)-Distribution (uniform / inverse-U-shaped), to model the effect of the hazard-rate function; (4) linear and quadratic terms for the Sequential effect, calculated as the difference between the current trial foreperiod and the previous trial foreperiod, such that positive values indicate the previous trial was longer than the current trial, and vice-versa for negative values; (5) The interaction terms between Foreperiod duration, Cue and FP-Distribution, and between Sequential effect, Cue and FP-Distribution. For simplicity, we assumed no interaction between sequential effect and foreperiod duration, e.g. we assumed that the cost in performance for a current trial of 900 ms and previous trial of 500 ms equals the cost of a 1300 and 900 ms pair of trials. To reduce computational complexity, all continuous factors were Z-scaled. To allow the computation of Sequential effects, the first trial of each session for each participant was discarded from analysis (total of 100 trials). Treatment contrasts coding scheme was used for Cue, with the uninformative condition set as the reference level, and sum contrasts coding scheme was used for FP-Distribution. Statistical significance for main effects and interactions was determined via a likelihood-ratio (LR) test against a reduced nested model excluding the fixed term (i.e. type-II sum of squares, SS). Statistical significance for parameter coefficients was determined according to Wald z-test (Fox, 2016).

In addition to the fixed effects, we considered the Z-scaled current trial number (i.e. the running trial identifier for the given session) as a covariate, in order to capture effects of fatigue and training along the experiment (Baayen and Milin, 2010). Since the different experimental groups may have experienced different fatigue or training effects, we additionally considered the interaction between FP-Distribution and trial number. Covariates were added to the model if the extended model converged and was found to significantly improve fit (*p* < .05) in an LR test against the model without the covariate (Bates et al., 2015a).

The model’s random effect structure was selected according to the model that was found to be most parsimonious with the data, i.e. the fullest model that the data permits while still converging with no singular estimates (Bates et al., 2015a), in order to balance between type-I error and statistical power (Matuschek et al., 2017). This was achieved by starting with a random intercept-by-subject-only model, and continuing to a model with random slopes for fixed terms by subject and their correlation parameters, and from there to a random interaction slopes by subject model, testing for model convergences in each step. Models that failed to converge were trimmed by the random slope with the least explained variance and were retested. Finally, we tested whether the model supports random slopes for the aforementioned covariates.

To provide support for null results (*p* < .05), we additionally modeled the data using a Bayesian GLMM, with weakly informative priors (Gelman et al., 2017) on the model’s fixed and random effects (*N*(0, 10)) and correlation (*LKJ*(2)) parameters, using the default mean for the intercept (298), and using informative shape parameters (*gamma*(0.02, 12.0)) according to Lo & Andrews (2015). Posterior distributions were constructed using four Markov chain Monte-Carlo (MCMC) chains and 20,000 iterations per chain, with the first 2,000 samples used as warmup. The large number of iterations was required in order to calculate a stable Bayes Factor (BF). BFs were calculated by comparing the marginal likelihood between the full model and a nested null model, with marginal likelihood estimated by 100 repetitions of bridge sampling (Gronau et al., 2017). BFs are reported with the null results in the nominator (*BF*_01_ or log *BF*_01_ for *BF*_01_ > 100), representing by how much the data is supported by the null model relative to the full model, along with range and the proportional estimation error (as in Morey & Rouder, 2018).

Analyses were performed in R v4.0.3 using R-studio v1.3.959 (R Core Team, 2018). Frequentist modeling was performed using the lme4 (Bates et al., 2015b) package, Bayesian modeling was performed using the brms package (Bürkner, 2017), and additional model diagnostics were performed using the performance package (Lüdecke et al., 2020). An R-markdown file describing all the model fitting steps and diagnostic checks on the final model is available at the project’s OSF repository (see Data Availability statement)

## Results

Reaction times (RTs) were modeled using a GLMM with FP-Distribution (uniform / inverse-U-shaped) as a between-subject fixed term and FP-Duration (continuous), Sequential effect (continuous), and Cue (valid / invalid / uninformative) as within-subject fixed terms, as well as the full interaction terms between FP-Duration, FP-Distribution, and Cue, and between Sequential effect, FP-Distribution, and Cue. Trial number and the interaction between trial number and FP-Distribution were added as covariates, and we allowed for a random intercept and a random slope for the linear term of FP-Duration and Cue by participant.

### Effects of foreperiod and spatial attention

Results showed that the *FP-RT slope*, the decrease in RT as foreperiod increases, changed with distribution, for each of the cues (see **Fig. 3**). We observed a significant main effect for FP-duration (*χ*^2^(2) = 864.59, *p* < .001), with negative linear and positive quadratic terms, consistent with the classic effect of foreperiod on RT and its expected shape, thought to reflect the increasing conditional probability along with the increase in the temporal uncertainty as the foreperiod duration becomes longer (Niemi and Näätänen, 1981). We additionally observed a main effect for Cue (*χ*^2^(2) = 19.90, *p* < .001), indicating the expected effect of spatial attention on RT. This effect was reflected by a large benefit in RT for valid vs. uninformative cues (*β* = −10.146, *t* = −12.582, *p* < .001) as well as a smaller but significant cost for invalid vs. uninformative trials (*β* = 2.666, *t* = 2.530, *p* = .011). Most importantly for the purpose of this study, we found no significant interaction between Cue and FP-Duration (*χ*^2^(4) = 5.862, *p* = .210), indicating that the effect for cue did not vary with foreperiod and supporting the hypothesis that spatial attention does not affect the FP-RT slope.

**Figure 3.**
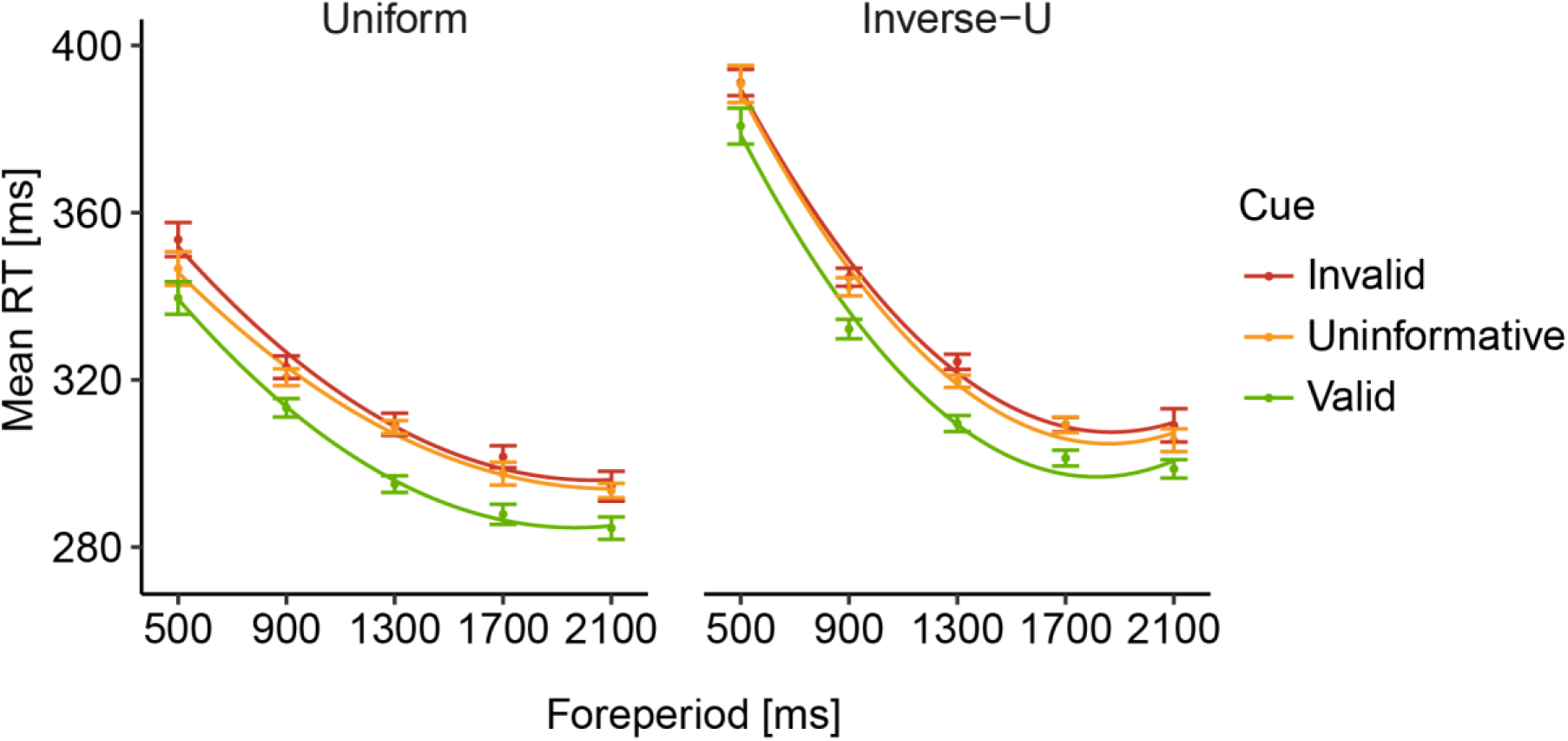
*Effect of hazard-rate function on RTs*. Mean reaction time (RT) for the uniform (left) and inverse-U-shaped (right) distributions. Each graph depicts group averaged mean reaction time (colored dots) with 2^nd^ degree polynomial fit (colored lines). Error bars represent ±1 standard error from the group mean, corrected to within-subject variability (Cousineau & O’Brien, 2014). *N*=20 for each group

### Effects of the hazard-rate function

The between-group variable of FP-Distribution (uniform / inverse-U-shaped) was used to assess the involvement of expectations based on the hazard-rate function on the foreperiod effect, and the relation of this effect to spatial attention. Findings showed no main group effect of FP-Distribution on RT (*χ*^2^(1) = 0.601, *p* = .435), indicating that both groups had similar overall RT. However, there was a significant interaction between FP-Distribution and FP-Duration (*χ*^2^(2) = 102.68, *p* < .001), indicating that, consistently with previous findings (e.g., Cravo et al., 2011; Trillenberg et al., 2000), the effect of foreperiod on RT was modulated by the prior distribution from which they originated, i.e. by their hazard-rate functions. Importantly for the goal of this study, there was no evidence that this effect of FP-Distribution on FP-Duration was modulated by the validity of the cue, as reflected by an insignificant interaction between Cue, FP-Distribution, and FP-Duration (*χ*^2^(4) = 4.699, *p* = .320). This suggests that the effect of the hazard-rate function on foreperiod, was independent of spatial attention. As expected, no significant interaction was found between Cue and FP-Distribution (*χ*^2^(2) = 0.050, *p* = .975).

### Sequential effects

To test for the existence of sequential effects, we calculated the difference between the FP-Duration of one trial and the FP-Duration of the previous trial (*FP*_*current*_ − *FP*_*previous*_). Consistently with previous studies (Alegria and Delhaye-Rembaux, 1975; Niemi and Näätänen, 1981), results showed an asymmetrical sequential effect on RTs, such that RTs were slower when the current trial was shorter than the previous trial (negative values in **Fig. 4**), but were not affected when the opposite was true (positive values in **Fig. 4**), leading to a quadratic relation with RT (*χ*^2^(2) = 1644.5, *p* < .001). The lack of effect when a trial is longer than its previous trial is thought to result from the combined contribution of sequential and hazard-rate effects: sequential effects erroneously guide expectations toward an early timing leading to lower performance; but, given that the target has not appeared at the earlier time, the conditional probability increases and expectation grow following the hazard rate function, leading to higher performance. Combined, the result is no enhancement or decrement of performance at late time points. Additionally, results revealed that this effect was significantly modulated by the FP-Distribution (*χ*^2^(2) = 28.924, *p* < .001), with linear component being more negative for the inverse-U compared to the uniform distribution. This finding, also consistent with previous findings (Niemi and Näätänen, 1981), supports the involvement of the hazard rate function in this effect. Generally, these findings demonstrate that expectations based on the hazard-rate function and sequential effects each had a unique contribution to the resulting RTs, along with a synergetic effect between them.

**Figure 4.**
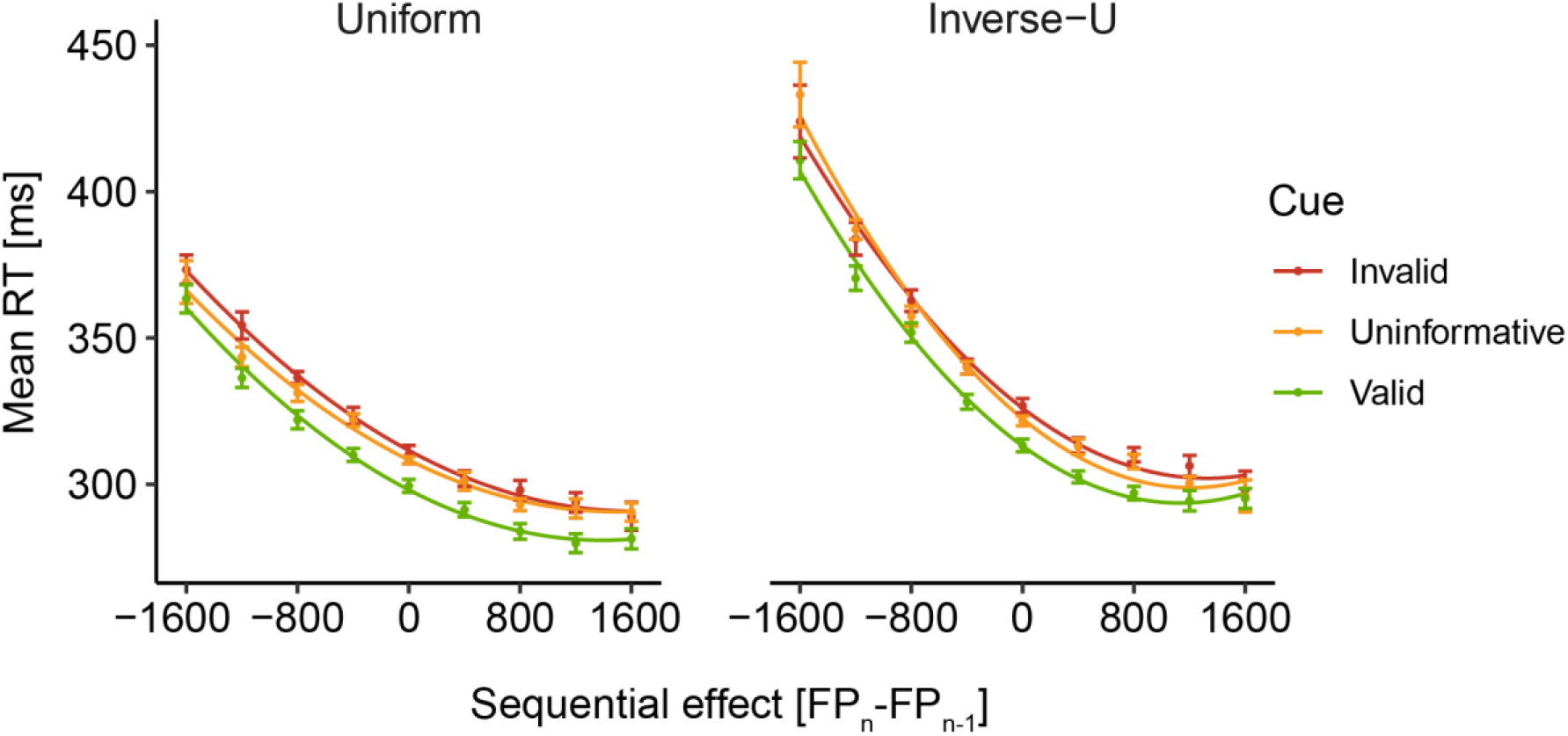
*Sequential effect on RTs*. Mean reaction time (RT) for the uniform (left) and inverse-U-shaped (right) distributions, with x-axis depicting the sequential effect (difference between current (FP_n_) and previous (FP_n−1_) trial foreperiod). Each graph depicts group averaged mean reaction time (colored dots) with 2^nd^ degree polynomial fit (colored lines). Error bars represent ±1 standard error from the group mean, corrected to within-subject variability (Cousineau & O’Brien, 2014). *N*=20 for each group.

We next tested whether these effects were modulated by spatial attention, by examining the interaction between them and Cue. Results showed no significant interaction between Sequential effect and Cue (*χ*^2^(2) = 1.177, *p* = .882), nor a significant three-way interaction between Sequential effect, FP-Distribution, and Cue (*χ*^2^(4) = 2.585, *p* = .630). Both results suggest that, as the hazard-rate effects, sequential effects are independent of the spatial locus of attention.

Model estimates for all fixed factors described are depicted in **Fig. 5**. Model estimates for covariates and additional model information can be found online in the project OSF repository (see Data Availability statement).

**Figure 5.**
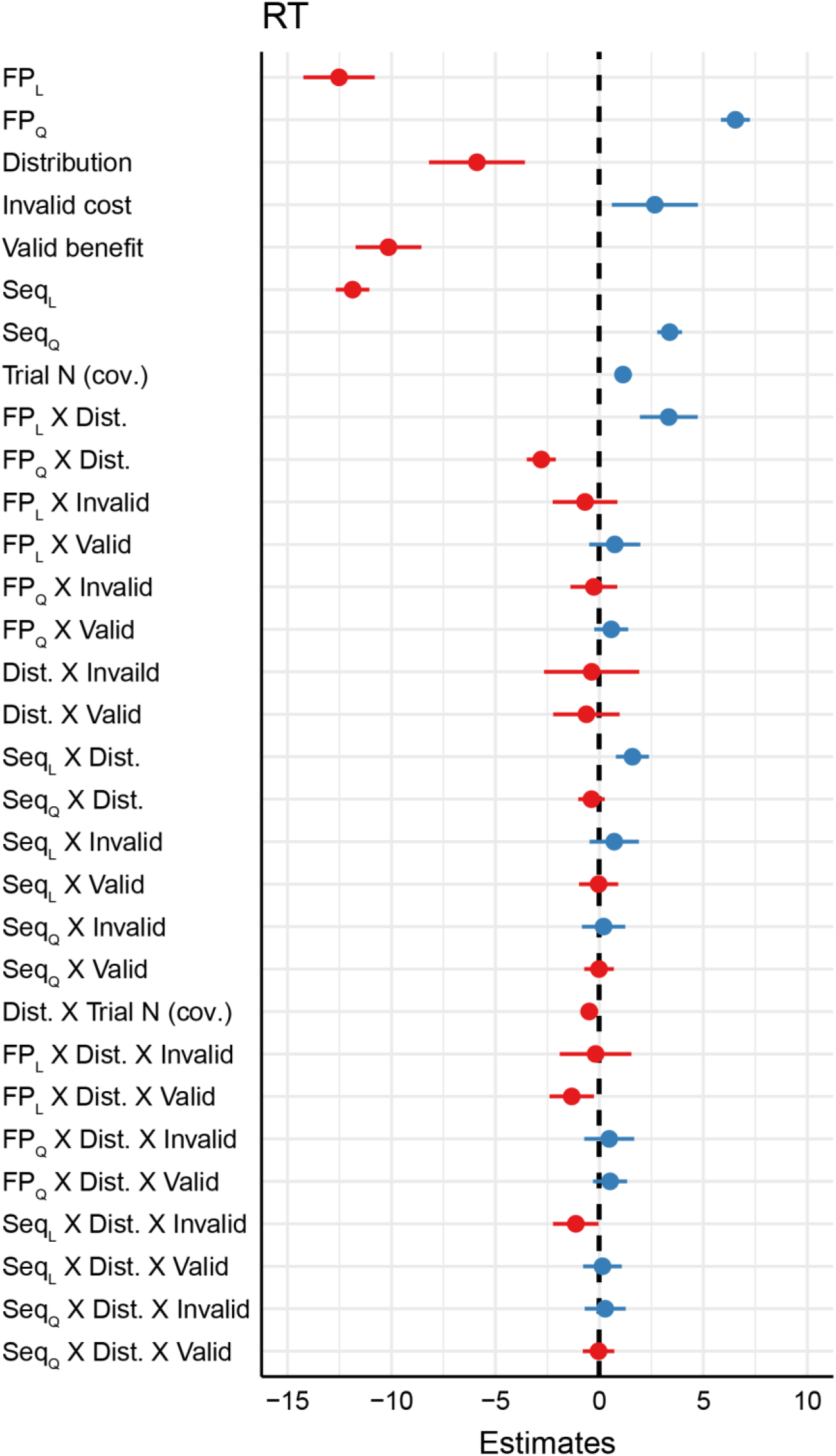
*Model estimates*. Forest plot of fixed factors estimates, modeled using a GLMM assuming a gamma response family and identity link function (estimates are given in ms units), and depicting mean in respect to the reference level (uninformative cue type). All continuous factors were scaled and centered. Positive values depicted in blue and negative values in red. Horizontal lines depict 95% Wald confidence intervals. Dashed vertical line centered at zero-sized estimate. Valid and invalid terms are relative to uninformative cue condition. *FP*_*L*_ = linear component of Foreperiod duration; *FP*_*Q*_ = quadratic component of Foreperiod duration; Dist = FP-Distribution; *Seq*_*L*_ = linear component of Sequential effect; *Seq*_*Q*_ = quadratic component of Sequential effect; Interaction terms denoted by X

### Bayesian modeling

Our results indicated that there was no evidence for a three-way interaction between Cue, FP-Distribution, and FP-Duration, as well as no three-way interaction between Cue, FP-Distribution, and Sequential effect. To examine whether the evidence supports these null results, we constructed a Bayesian GLMM using the same model terms. Model estimates closely resembled the coefficients found in the frequentists model. We compared the resulting Bayesian model with two nested models, each lacking the corresponding three-way interaction term. Results showed large support for the null model lacking the FP duration three-way interaction term compared to the full model (mean log *BF*_01_ = 8.483 ± 0.002%, range 8.289-8.681), and similarly large support was observed for the null model lacking the Sequential effect three-way interaction term compared with the full model (mean log *BF*_01_ = 9.969 ± < .001%, range 9.731-10.146). Both results support the conclusion that temporal expectations based on hazard-rate function and sequential effects are independent of spatial attention. Additional modeling information can be found online (see Data Availability statement).

## Discussion

In this study, we examined whether the spatiotemporal model which was proposed to account for cue-based temporal expectation also carries for temporal expectation based on contextual information, i.e. the hazard-rate function and sequential effects. By varying the foreperiod, we observed the established FP-RT slope effect, with RT decreasing as foreperiod increases. This FP-RT slope changed according to the hazard-rate function, which was manipulated by varying the foreperiod distribution. In addition, we found the expected asymmetrical sequential effect: slower RTs for trials in which the foreperiod was *longer* than their previous trial, and no opposite effect for trials in which the foreperiod was *shorter* than the previous trial. Critically, all these effects were unaffected by spatial attention – similar modulations of expectations were found in both attended and unattended spatial locations. This indicates that temporal expectations based on contextual information – the hazard-rate function and recent previous experiences – are independent of spatial attention.

### The spatiotemporal model of temporal expectation

Doherty et al. (2005) were the first to demonstrate an interaction between cue-based temporal and spatial attention in early visual event-related potentials (ERP) components. They presented participants with moving objects that disappeared behind an occluder and reappeared in an expected or unexpected location and/or time. Participants were requested to indicate whether a target dot was presented on the reappearing object. Findings showed that when a target appeared at an expected location, the early visual P1 component was increased relative to an unexpected location, and this effect was enhanced when the target also appeared at the expected time. However, when a target appeared at the expected time but not the expected location, there was no enhancement relative to its appearance at an unexpected time and location, suggesting that early perceptual benefits of temporal attention depend on the allocation of spatial attention. This spatiotemporal synergism was not found in later ERP components, such as the P3, considered to be less affected by perceptual processes and more by response requirements, and also not in RTs.

In a later study by Rohenkohl et al. (2014), symbolic spatial and/or temporal cues predicted 80% validity the time and location of a grating-patch target, for which participants were requested to perform a non-speeded orientation discrimination task. Findings showed that valid temporal cues improved both RT and perceptual sensitivity relative to invalid cues, but that this effect was limited to trials where spatial attention was focused at the target’s location. These findings provided, again, evidence for a strong synergistic interaction between temporal and spatial expectations in a discrimination task. Consistently, recent evidence by Seibold et al., (2020) showed that temporal attention boosts the effect of spatial attention on early ERP components in a visual search task.

This evidence of a tight link between spatial attention and cue-based temporal expectation led Nobre and van Ede (2018) to propose their spatiotemporal neurophysiological model, which can account for these findings. According to this model, the interaction between spatial and temporal processes stems from time-specific synchronization of spatially-specific neural populations at the attended retinotopic receptive-fields. These neurons, coding the attended location and relevant features, acquire a temporal structure from repeated exposure to the temporal cues, which affects them but not populations outside the receptive-field (Nobre and van Ede, 2018). This model was developed based on evidence on cue-based expectation but was never before examined for other sources of temporal expectations. The present evidence indicates that hazard-rate and sequential effects do not depend on spatial attention, suggesting that these forms of expectation cannot be explained by the spatiotemporal mechanism proposed by Nobre and van Ede. This further suggests that cue-based temporal expectation and temporal expectation that are driven by contextual information, which are often described as two manifestations of the same expectation process, likely rely on distinct neural mechanisms. This evidence is consistent with studies that dissociated hazard rate effects and cue-based temporal expectation and found that these two sources of expectations share some, but not all, of their underlying brain networks (Lima et al., 2011; Coull et al., 2016; Amit et al., 2019). More generally, this conclusion is compatible with the increasing recognition in this field that there is no single unified expectation mechanism, but that distinct sources of temporal expectations facilitate performance via distinct neural mechanisms (van Ede et al., 2020).

### Spatiotemporal synergism and cue-based expectations

It is important to note, however, that evidence regarding the dependency, or lack thereof, of cue-based temporal expectation on spatial attention, is ambivalent. In addition to the supporting evidence described above, a few studies provided evidence challenging this interaction. For example, MacKay & Juola (2007) used a visual search task in a rapid stimulus visual presentation (RSVP) stream of letters. Visual cues were provided to indicate the time, location, or both of the target letters, and a discriminate task was performed on the cued targets. Findings showed that both types of cues were effective on their own and their combined effect was additive, indicating that there was no interaction between temporal and spatial attention. In a later study, Weinbach et al. (2015) used a spatiotemporal cueing paradigm and showed that temporal cueing improves RT even when coupled with an invalid spatial cue. Moreover, there was no interaction between the effect of the temporal and the spatial cues, indicating that enhancement resulting from temporal attention was not affected by spatial attention. The authors noted that the discrepancy between their findings and previous findings could have stemmed from differences in task demands: whereas most previous studies used demanding perceptual discrimination tasks, they used a speeded-RT detection task. Another study by Rolke et al (2016) investigated the combined influence of temporal, spatial, and feature-based attention and found no synergetic effects between spatial and temporal attention when spatial attention was manipulated. In that study, temporal expectations were manipulated implicitly, whereas spatial attention was manipulated explicitly using symbolic attentional cues. Findings showed no spatiotemporal interaction, and therefore it was suggested that this interaction occurs only when attention is manipulated similarly in both modalities (Seibold et al., 2020).

### Temporal attention and temporal expectation

The apparent discrepancies among different findings on spatiotemporal dependency could be accounted for by the dissociation between attention and expectation processes. According to one view, described in Summerfield & Egner (2009), expectation reflects the narrowing down of the probability space of possibilities, constructed according to prior knowledge; whereas attention is the selection of specific, goal-relevant information that should be prioritized. Both attention and expectation coexist and are often entangled – e.g., cueing to the left visual field increases our expectation of encountering a target at that location, and induces a shift of attention that prioritizes information on that particular visual space. Tailored experimental designs can dissociate attention and expectation, as was demonstrated in visual spatial attention and feature attention studies (Summerfield and Egner, 2009, 2016; Kok et al., 2012).

Similar to spatial cues, temporal cueing paradigms often create a symbolic association between a certain cue and a specific target onset time. Thus, the onset of the cue induces an attentional shift which prioritizes information processing around the cued time interval. In addition, in these designs, the repeated exposure to target onset after a cue changes the probability space and induces *temporal expectation,* which is independent of attention according to the definition described above (Summerfield & Egner, 2009; but see Nobre & van Ede (2018) for a different approach). Therefore, according to this view, in these designs, temporal attention often coincides with temporal expectation, although specific experimental designs can dissociate these functions (Denison et al., 2019, 2021). Importantly, according to this definition, both hazard-rate function and sequential effects can be viewed as forms of temporal expectation, as they narrow down the probability space.

We hypothesize that this proposed dissociation between expectation and attention could account for the discrepancies between previous studies on the spatiotemporal dependency, with temporal attention being spatially-specific, while temporal expectation remaining independent of the spatial locus of attention. This, in turn, could explain the results observed here – since the hazard rate and sequential manipulations affect only temporal expectation and not attention, their manifestations were free of spatial constraints.

### Conclusions

This study examines the relation between spatial attention and two forms of temporal expectation – those based on the hazard-rate function, the moment-by-moment increase in a target’s conditional probability over time, and those based on sequential effect. Our results showed that both forms of temporal expectations are independent of spatial attention. We conclude that the benefit from these forms of expectation is not spatially-specific, but rather reflects a general non-specific enhancement that is not accompanied by shifts of attention. Furthermore, we suggest that the spatiotemporal neurophysiological model proposed by Nobre and van Ede (2018) to explain cue-based expectation cannot account for hazard-rate and sequential expectation effects. Future studies are encouraged to examine the dissociation between different mechanisms of temporal expectation, and to refine the terminology to reflect this dissociation.

## Acknowledgements

We thank Danielle Allon and Keren Nistor for their assistance in running the experiment. This study was funded by the Israel Science Foundation grant 1960/19 to S.Y-G.

## Data availability

The datasets generated by this study and an R-markdown file that reproduces all the reported modeling, statistical analyses and graphs within the paper are uploaded to the Open Science Foundation repository and are available at: https://osf.io/25gzj

## Notes

*Conflict of interest*. We have no conflict of interest to disclose.

https://osf.io/25gzj/

### Competing Interest Statement

The authors have declared no competing interest.

